# The organization of individually mapped structural and functional semantic networks in aging adults

**DOI:** 10.1101/2021.09.29.462441

**Authors:** W. Tyler Ketchabaw, Andrew T. DeMarco, Sachi Paul, Elizabeth Dvorak, Candace van der Stelt, Peter E. Turkeltaub

**Affiliations:** Center for Brain Plasticity & Recovery, Georgetown University Medical Center; Research Division, National Rehabilitation Hospital

## Abstract

Language function in the brain, once thought to be highly localized, is now appreciated as relying on a connected but distributed network. The semantic system is of particular interest in the language domain because of its hypothesized integration of information across multiple cortical regions. Previous work in healthy individuals has focused on group-level functional connectivity (FC) analyses of the semantic system, which may obscure interindividual differences driving variance in performance. These studies also overlook the contributions of white matter networks to semantic function. Here, we identified semantic network nodes with a semantic decision fMRI task in 53 typically-aging adults, characterized network organization using structural connectivity (SC), and quantified the segregation and integration of the network using FC. Hub regions were identified in left inferior frontal gyrus. The individualized semantic network was composed of three interacting modules: 1) default-mode module characterized by bilateral medial prefrontal and posterior cingulate regions and also including right-hemisphere homotopes of language regions; 2) left frontal module extending dorsally from inferior frontal gyrus to pre-motor area; and 3) left temporoparietal module extending from temporal pole to inferior parietal lobule. FC within Module3 and integration of the entire network related to a semantic verbal fluency task, but not a matched phonological task. These results support and extend the tri-network semantic model (Xu et al., 2017) and the controlled semantic cognition model (Chiou et al., 2018) of semantic function.

## Introduction

### Semantic function in the brain

Over the past decade, the study of the neurobiology of language has undergone a paradigm shift. Early neuroimaging studies of language focused on localization of function to specific brain areas, but more recent work has taken a network-based perspective that considers information transfer between neural processors (Medaglia et al., 2015). The semantic network, hypothesized to function in a way that binds lexical items to representations from distributed brain regions (Chiou et al., 2018), is of particular interest for elucidating how behavior is supported by the organization and interaction of relevant brain networks. Previous neuroimaging studies of semantic function in the brain have found group-level activation in distributed regions, including canonical language regions in left inferior frontal gyrus (IFG) and left temporoparietal cortex, in addition to “domain general” regions like bilateral medial prefrontal cortex that are not specific to language function (Braga et al., 2020; Fedorenko & Thompson-Schill, 2014).

From a network perspective, functional connectivity studies have been cited as support for two theories of semantic function. First, it has been proposed that semantic function is coordinated by a highly influential “hub” region in anterior temporal lobe (ATL) interacting with peripheral “spokes” to form polymodal representations which are linked to lexical items (Patterson & Ralph, 2016). An expansion of this hub-and-spoke model, the “controlled semantic cognition” (CSC) model (Chiou & Lambon Ralph, 2019), broadens the concept of a unitary central hub to instead cover a graded span of cortex in ATL. The CSC model additionally includes left inferior frontal regions which are thought to play a role in selecting the proper representation from semantically-related competitors (Chiou et al., 2018). Alternatively, a tri-network model of semantic functioning has been proposed, consisting of a frontoparietal network, a perisylvian network, and parts of the default mode network (Xu et al., 2017). Under this model, semantic representations are accessed through perisylvian and default mode networks, with frontoparietal network performing roles in cognitive control.

However, the evidence underpinning these theories is lacking in two respects. First, these studies have not accounted for interindividual variability in the arrangement of the language network (Fedorenko et al., 2010). The definition of the semantic network in prior studies was driven by group-level activation on fMRI tasks, which may be overlooking idiosyncratic features of individuals’ brain network organization (Gordon et al., 2017). A high degree of interindividual variability has been reported in the language domain in particular (Fedorenko & Thompson-Schill, 2014), and individually defining language regions based on first-level fMRI activation may be important for understanding individual differences in language abilities (Nieto-Castañón & Fedorenko, 2012). Secondly, prior work studying the semantic network in healthy individuals has primarily focused on functional connectivity studies. Such studies do not consider the presence of white matter networks which present neurophysiological and anatomical constraints on connectivity (Chiou et al., 2018; Friederici, 2015). To address these gaps, we localize the semantic network at the individual level (Nieto-Castañón & Fedorenko, 2012) using a validated semantic decision fMRI task (Wilson et al., 2018), then use structural connectivity to inform functional connectivity analyses of these individualized semantic networks (ISNs) using graph theory. Finally, we relate network properties to performance on semantic and phonological fluency tasks.

### Graph theory and small-world brain networks

The brain has an intrinsic network organization (Fox et al., 2005) comprised of many local connections between neighboring regions as well as sparser longer-distance connections bridging these communities (Meunier et al., 2010; Sporns & Betzel, 2016). Graph theory, a field of mathematics that quantifies relationships between objects in a set, defines networks with these characteristics as “small-world” (**Figure S1**), and previous work suggests this is the most salient feature of brain network organization (Bassett & Bullmore, 2006; He & Evans, 2010). In graph theory analyses, networks are generally conceptualized as gray matter processors (“nodes”) linked together by white matter fibers, forming connections or “edges” (Bullmore & Sporns, 2009). Network properties can be summarized by segregation, i.e., connectivity within communities of nodes, and integration, i.e., connectivity across communities (Rubinov & Sporns, 2010; Sporns, 2013). Segregation is often summarized for an entire network with clustering, the fraction of a node’s connected neighbors that are also connected neighbors (**Figure S1**) (Rubinov & Sporns, 2010). Integration is often summarized with shortest path length, the sum of edges that constitute the most direct route between a node pair, or with global efficiency, the inverse of shortest path length.

As an alternative to whole-network measures, segregation and integration can be examined by comparing the connectivity of algorithmically-detected communities of nodes, also called modules, that are more strongly connected within modules than across modules (Sporns & Betzel, 2016). Module-based analyses can therefore describe contributions of specific groups of nodes within a network, as compared to summary measures like clustering and global efficiency which describe the entire network. Modules are critical features of brain networks, detectable at both small and large scales (Bassett & Bullmore, 2017; Sporns & Betzel, 2016), and disruption of the brain’s modular arrangement is associated with deficits across cognitive domains in stroke (Siegel et al., 2016).

In addition to properties of whole networks and subnetworks (modules), graph theory also quantifies contributions of individual nodes and edges to the network at large. In describing the influence of single regions, graph theory analyses often report disproportionately-strongly connected “hub” nodes (van den Heuvel & Sporns, 2013b). These hubs are thought to play important roles in coordinating information flow in brain networks. Hubs can also be viewed as part of the modular framework; a distinction is made between hubs that are highly connected within a module (“provincial” hubs) and hubs that are highly connected across modules (“connector” hubs) (Power et al., 2013). Evidence from lesion studies suggests that connector hubs are more important for cognitive function(Gratton et al., 2012), but the relative functional relevance of provincial and connector hubs in the brain is still under debate (van den Heuvel & Sporns, 2013b).

### Studying brain networks with functional and structural connectivity

Nodes are usually defined using a whole-brain parcellation or by selecting regions of interest (ROIs) from an atlas, most often based on group-level activation on an fMRI task. Edges in brain networks are then defined as the connectivity between node pairs; for fMRI studies, this is most often either structural or functional connectivity (Wang et al., 2015). Diffusion tensor imaging (DTI) is a common structural connectivity (SC) approach capable of investigating anatomical white matter networks, thereby elucidating anatomical routes through a network, but agnostic to whether or not information is actually being exchanged along these routes (Sporns, 2013). Conversely, functional connectivity (FC) calculates correlations in fMRI BOLD activity over time between gray matter regions, often in the absence of a task. Nodes connected by an edge with strong FC are considered to be communicating as part of a functional network, without regard for the proximity of the two regions or the presence of white matter (anatomical) connections between them (Sporns, 2013). SC and FC are closely related, but the relationship is incompletely understood and there is no direct one-to-one mapping between the two (Goñi et al., 2014; Wang et al., 2015). There is strong evidence that more direct structural connections (i.e., shorter path length) are associated with stronger FC (Goñi et al., 2014; Honey et al., 2009), although this does not explain why some region pairs without direct structural connection report consistently high FC across individuals (Honey et al., 2009; Wang et al., 2015). Still, the relationships between direct structural connection and stronger FC suggest that structural networks represent an anatomical substrate upon which functional networks are built (van den Heuvel & Sporns, 2013a). Therefore, under this framework, we use SC in the present study to reveal anatomical routes within the semantic network and FC to investigate how information actually flows through individually-defined semantic networks in intact brains.

In this study, we use the segregation, integration, and influence framework (Rubinov & Sporns, 2010) to describe the organization of the individually-defined semantic network in typically-aging adults. Understanding the typical organization of the semantic network in this age group will be essential to studying how stroke or neurodegeneration disrupts the network, and how this disruption relates to semantic deficits in aphasia (Turkeltaub, 2019). We define the functional semantic network in each individual using an fMRI task to account for interindividual variability. Because the organization of these networks is constrained by anatomical connections, we use SC to identify hub regions and interrogate the modular arrangement of the individualized semantic network and FC to investigate segregation and integration properties. Finally, we relate these FC measures to performance on semantic and phonological fluency tasks.

## Methods

### Participants

Participants were 53 native English speaking, typically-aging adults (29 female, mean age 60.1 ± 11.9 years) with no history of significant neurological or psychiatric condition. This group was recruited as a matched control sample for a study of stroke survivors, so participants who had medical conditions that might affect brain health such as hypertension and diabetes mellitus type 2 were not excluded. All participants scored in the typical range for their age and education on the Montreal Cognitive Assessment based on published norms (Rossetti et al., 2011). The research protocol was approved by the Georgetown University Medical Center Institutional Review Board, and written informed consent was acquired for all participants before enrollment in the study.

### Image acquisition and preprocessing

#### Functional imaging

Both fMRI task and naturalistic resting-state scans were acquired with the same parameters: A BOLD T2^*^-weighted scan (TR = 794ms, 48 2.6mm slices with 10% gap, 2.9mm voxels, FOV = 211mm, matrix = 74×74, FA = 50°, SMS = 4), consisting of 504 volumes lasting 6:40 for the task and 1024 volumes lasting 14:37 for the naturalistic resting-state. A common preprocessing pipeline was run on both task and resting-state scans in AFNI (Cox, 1996), including realignment for head motion, despiking, smoothing with a 6mm FWHM kernel, temporal high-pass filtering at 0.01 Hz, and detrending. Images were coregistered to an MPRAGE which was parcellated using the structural connectivity-derived Lausanne atlas (Hagmann et al., 2008) at scale125 (234 nodes).

#### Task

The fMRI task was a validated adaptive semantic decision task, described in detail in (Wilson et al., 2018). Briefly, in the language condition, participants decide if two printed words are semantically related. In the control condition, participants decide if two false font strings are matching (**Figure 1A**). Responses to each trial were collected via button press. The participants complete 20 alternating blocks of language and control conditions, with 4-10 items in each block depending on adaptive difficulty level. To maintain similar levels of effort across different levels of language ability, a staircase procedure adapts item difficulty to each individual participant’s performance. Statistical analysis was conducted using a whole-brain GLM was estimated using the *fmrilm* function from FMRISTAT (Worsley et al., 2002), with covariates including the time-course of a white matter and CSF seed, and the six head-motion parameters not convolved with the HRF. The semantic decision task was modeled using two alternating boxcar functions (corresponding to the language and control conditions), convolved with the HRF. The contrast of interest was semantic greater than control condition, specified as [1 -1].

**Figure 1.**
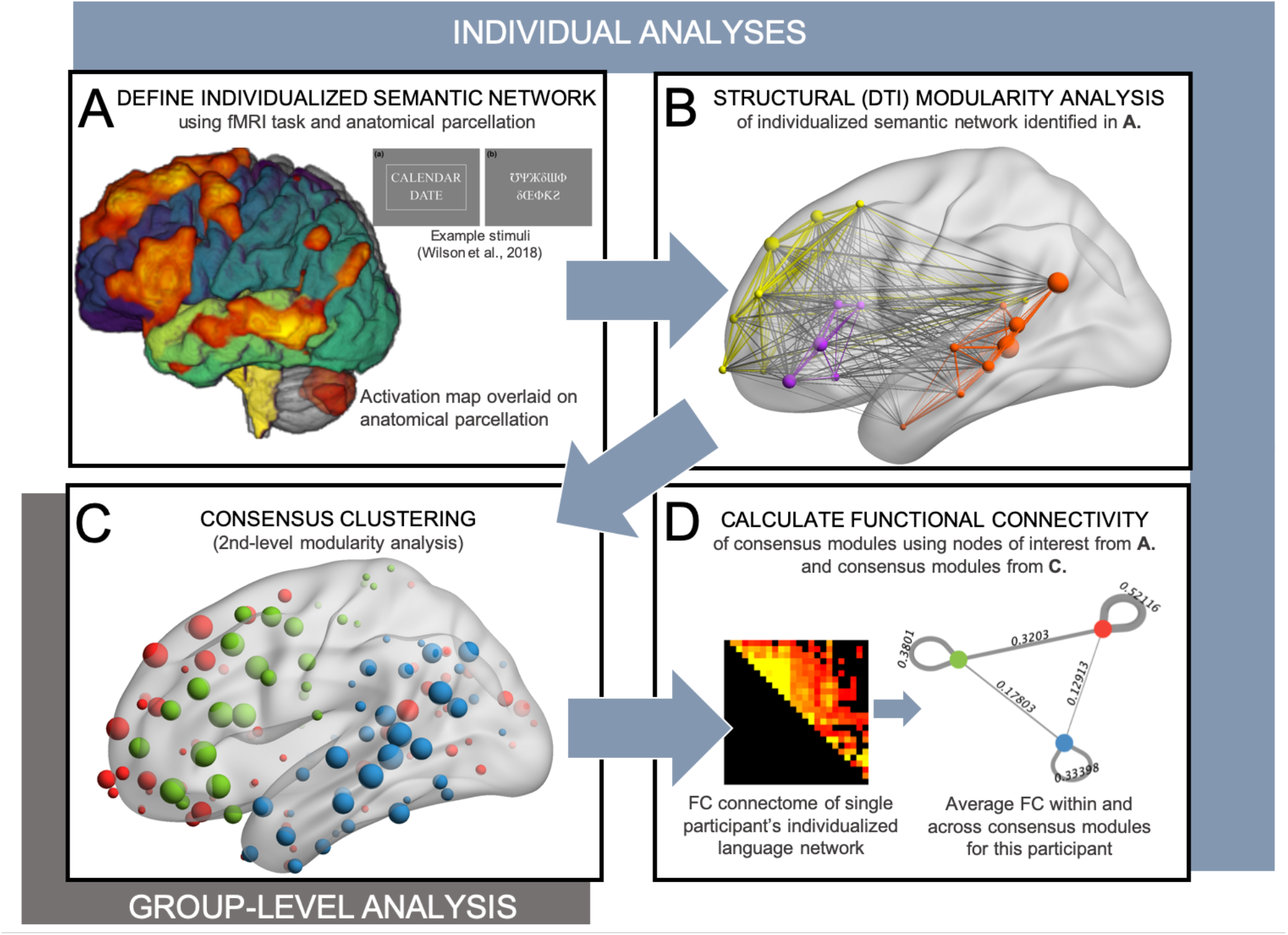
Graphical summary of the imaging and connectivity analyses. **A**. Individualized semantic networks (ISNs) were localized using a Semantic Decision fMRI task (Wilson et al., 2018). **B**. First-level modularity analysis was then performed using structural connectivity data (DTI) to investigate community structure of each participant’s ISN. **C**. To enable comparisons across ISNs, consensus clustering (Lancichinetti et al., 2012) was used, assigning each node to a consensus module. **D**. Functional connectivity was then calculated within and between each of the consensus modules in each participant’s ISN.

#### Derivation of individualized semantic networks

After coregistering the resulting T map to the parcellated native-space MPRAGE, mean T scores were calculated for each of the 234 nodes, and the top 10% (24 nodes) were used to define an individualized semantic network (ISN) for each participant. This top-10% cutoff was chosen rather than a p-value cutoff in order to generate ISNs with the same number of nodes in each individual, sized similarly to previously-established resting-state networks (Gordon et al., 2016; Yeo et al., 2011). At higher cutoff values (15% and higher), ISNs contained many nodes that do not activate with the task (mean activation T < 1), and at lower cutoff values the networks were not fully connected (i.e., some nodes were not connected to any other nodes). The cutoff of top 10% activation was therefore deemed appropriate for these analyses. Results were similar at top 11% and 12% cutoffs (data not shown) suggesting the cutoff did not drive our results.

#### Naturalistic resting-state paradigm

Participants watched a 14.5-minute movie excerpt in the scanner during the naturalistic resting-state scan. The excerpt comes from *The Gruffalo* (2009), starting just after the opening credits (1m29s mark) and ending on a shot of the Gruffalo’s face 16 minutes into the film. Prior work indicates that naturalistic movie watching induces the same patterns of connectivity as normal resting state paradigms with higher reliability and less head motion (Vanderwal et al., 2015) and may be helpful for elucidating individual differences (Vanderwal et al., 2017). After the common preprocessing steps described above, the following nuisance regressors were removed before whitening the data to give a final residual file: signal from white matter, CSF, scalp, and skull as well as head motion and its first temporal derivates. Functional connectomes were generated for each individual by extracting the average BOLD timecourse from each node of the anatomical parcellation and calculating the correlation (Pearson’s r transformed by Fisher Z) between timecourses from each pair of nodes in that participant’s ISN.

#### DTI acquisition and pre-processing

To quantify white matter connections, multi-shell High Angular Resolution Diffusion Imaging scans were acquired (HARDI; TR = 4.7 or 5.0s, TE = 0.082 s, readout time = 0.061 s, diffusion-weighted gradients: 81 directions at b= 3000, 40 at b= 1200, 7 at b= 0, 70 slices, 2 mm cubic voxels, flip angle = 90, phase encoding direction = anterior to posterior, partial fourier = 6/8, FOV = 232 mm, Matrix = 116 × 116, slice acceleration = 1). 6 volumes of reverse phase-encoded b=0 images were also acquired for susceptibility field estimation. Structural connectomes were constructed from the HARDI data through the MRtrix 3.0 software (Tournier et al., 2019) as described in full in (Dickens et al., 2021). Briefly, preprocessing followed the standard stepwise application of gaussian noise removal, gibbs ringing artifact removal, correction of distortions induced by motion, eddy currents, and magnetic susceptibility, and correction of B0 field inhomogeneity. Voxelwise fiber orientation distributions were computed using multi-shell, multi-tissue constrained spherical deconvolution (Jeurissen et al., 2014). The preprocessed diffusion-weighted images were upsampled to 1.3 mm prior to spherical deconvolution to increase anatomical contrast. Structural connectivity was quantified through 15 million streamlines generated by probabilistic anatomically-constrained tractography (Smith et al., 2012) on the normalized white matter fiber orientation distributions. Streamline density was made proportional to the voxel- and orientation-wise apparent fiber density through spherical-deconvolution informed filtering of tractograms 2 (SIFT2) (Smith et al., 2015). Specifically, edges of the structural connectome were generated by assigning streamlines (Daducci et al., 2012) to parcels of the Lausanne atlas at scale 125, and then multiplying each streamline by its respective cross-sectional multiplier derived by *tcksift2*. Overall, each edge value is directly proportional to the cross-sectional area of white matter connecting the two parcels.

### Graph theory analyses

#### Structural hub analysis

One prominent feature of brain networks is the presence of disproportionately highly-connected nodes, called hub nodes, which are thought to be important for relaying and coordinating information transfer within and between brain networks (van den Heuvel & Sporns, 2013b). The present study focused on structural hubs rather than functional hubs because FC doesn’t require anatomical connections, a theoretical criterion for network hubs based on neurophysiological constraints (Chiou et al., 2018; van den Heuvel & Sporns, 2013b). Further, based on lesion studies, structural hubs seem to be more relevant to cognitive function than functional hubs (Gleichgerrcht et al., 2015; Griffis et al., 2019; Reber et al., 2021). For each node in each ISN, the following graph measures were calculated (Sporns et al., 2007): strength, the sum of all connection weights of a given node; betweenness centrality, the number of shortest path lengths in the network that pass through a given node; and participation coefficient, a ratio of cross-to within-module connectivity for a given node. To find nodes with much stronger connectivity than the rest of the network, strength and betweenness were Z-scored within each individual ISN, and hub nodes were defined as nodes with Z > 1 for both metrics (Sporns et al., 2007). Participation coefficient (P), a measure of intermodular connectivity, was then used to characterize the hubs as either provincial (within-module) or connector (cross-module). Hubs with P < 0.3 were considered provincial hubs and hubs with P ≥ 0.3 were considered connector hubs (Power et al., 2013).

#### Structural connectivity modularity analysis

Brain networks are organized into communities, or modules, that are more strongly connected within a community than across communities (Meunier et al., 2010). Disruption of the modular arrangement of brain networks in stroke survivors is associated with cognitive deficits including aphasia (Siegel et al., 2016), and FC studies suggest the semantic network is highly modular (Yu et al., 2018). The present study uses SC to detect modules for two reasons. First, brain networks are constrained by anatomical connections. Secondly, this approach enables the examination of FC within and between modules without being biased by the community detection algorithm. That is, the community detection algorithm maximizes connectivity within modules, so the use of one modality (SC) to define the network permits the calculation of intra- and intermodular connectivity with a separate, complementary modality (FC) without so-called “double dipping.” Community detection was performed on SC data of each participant’s ISN using the Louvain algorithm with optimization (Que et al., 2015; Rubinov & Sporns, 2011), yielding anatomical subnetworks through which information can route in each individual (**Figure 1B**). Because semantic networks are individualized for these analyses based on fMRI task activation, the ISNs do not necessarily contain the same nodes across participants. To account for these differences, we derived consensus modules by finding group overlap in individual (first-level) modules. Specifically, consensus modules were derived using an adaptation of the consensus clustering procedure (Lancichinetti & Fortunato, 2012; Rasero et al., 2017). Briefly, an association matrix was constructed, counting the number of times across individuals that any two nodes occurred in the same module, and then the Louvain algorithm was run on this association matrix (**Figure 1C**), yielding consensus clusters of nodes that served as stable group-level modules. Each node was then assigned to a group-level module according to the consensus cluster analysis to permit a comparison of modules across participants’ ISNs.

#### Functional connectivity analyses

Once hub regions and consensus modules were derived using the SC data, FC was used to characterize how information is communicated through these anatomical subnetworks. To quantify segregation and integration in the whole network, three commonly-reported graph theory summary measures (Bullmore & Sporns, 2009) (**Figure S1**), which do not consider the presence of modules, were calculated using Brain Connectivity Toolbox (Rubinov & Sporns, 2010): 1) clustering, a measure of segregation; 2) global efficiency, a measure of integration; and 3) small world propensity, a combined measure that compares the observed clustering and global efficiency to values in ideal small-world networks. Small-world propensity contextualizes clustering and global efficiency by normalizing relative to an “ideal” small-world network (Muldoon et al., 2016). Ideal networks in this context are clustered similarly to a lattice graph (i.e., when every node is connected to all of its neighbors) and are similarly efficient to a random graph (i.e., when nodes have equal probability of being connected to any other node in the graph). For the consensus modules, segregation and integration was calculated (**Figure 1D**). Segregation was calculated as the average FC between all node pairs within each consensus module. Integration was calculated for each pair of consensus modules by taking the average connectivity of each node in one module to every node in the other module, i.e., cross-network connectivity. By calculating segregation and integration both at the whole-network and the modular level, we are able to interrogate the contributions to behavior of both the network-wide organization as well as the role of specific modules.

#### Behavioral testing

Verbal fluency is a cognitive ability involving the retrieval and articulation of information from memory (Shao et al., 2014). We examined verbal fluency for two reasons: (1) because matched semantic and non-semantic versions are available allowing us to examine the specificity of the network-behavior relationships as determined by our regressions and (2) because there is no ceiling effect on performance of verbal fluency tasks so there is sufficient variability for performing regressions.

##### Category [semantic] Fluency

participants were presented with a category and asked to list as many related words aloud as possible in 60 seconds. All participants performed two versions of the task, using the categories “animals” and “fruits & vegetables.” If a participant listed both a supraordinate item (e.g., “bird”) and accompanying subordinate items (e.g., “eagle” and “falcon”) only the subordinate items were counted.

##### Letter [phonological] Fluency

similarly, participants were presented with a letter and asked to list as many words as possible in 60 seconds that begin with that letter. All participants performed three versions of the task, using the letters *F, A*, and *S*. Proper names of people and places were not counted. If a participant listed two words related by morphology, derivations were counted as separate words (e.g., “farm” and “farmer”) but inflections (e.g., “float” and “floating”) were not. If the participant began listing numbers (“forty, forty-one”), they were redirected to give different words and only the first number was counted.

#### Backwards-elimination regressions with behavior

To test the hypothesis that language abilities are related to segregation and integration of the functional language network, backward-elimination linear regressions were performed in SPSS using verbal fluency scores as the dependent variable and measures of segregation and integration as predictors. Two separate models were run for each fluency task with main effects of the following predictors: 1) graph theory measures of the entire network (clustering and global efficiency); and 2) within- and cross-module connectivity of each of the 3 modules. All models included age and years of education as covariates. Model #1 therefore starts with 4 variables, clustering, efficiency, age, and education. Model #2 starts with 8 variables, connectivity within each of the three modules, connectivity across the three pairs of modules, and age and education. The backwards elimination procedure starts with main effects of all terms in the model and iteratively removes one variable with the highest (least significant) F-test probability, leaving at the end either a subset of the original predictors that contribute significantly or an empty model (to remain in the model, variables must have *P* < 0.10). The backward-elimination approach was selected because it considers all variables initially and is less susceptible to multicollinearity across predictors compared to other stepwise regression techniques (Myers et al., 2016).

## Results

### Semantic network localization

The fMRI task was successful in localizing language processors in each individual, identifying expected areas of consistent activation in left inferior frontal nodes and temporoparietal nodes, as well as domain-general processors in medial prefrontal cortex (**Figure 2A & B**). Our first analysis aimed to inspect the amount of interindividual variability in task activation patterns. This is important for establishing the utility of using individualized networks for our analyses. To quantify the consistency of activation patterns across individuals, each participant’s activation at every node was made into a 234-element vector, and correlations (Pearson’s r) were calculated on these nodewise activation vectors, computing correlations for each pair of participants. A summary of each participant’s similarity to the rest of the group was derived by calculating the average of their correlation to every other participant in the study (**Figure 2C**). The average of this summary similarity score across individuals was R^2^ = .514. This correlation analysis also revealed 2 outliers with r < .5 with the rest of the group. Upon visual inspection of their activation patterns, these two participants were found to be right-lateralized for language, and were excluded from all further analyses (**Figure 2B**, first 2 columns). After removing these two individuals, the average group correlation was R^2^ = .523. Only 1 node (left pars triangularis) was identified as part of the language network in all participants, and three others (left superior temporal sulcus, left pars opercularis, and left pars orbitalis) were identified in >95% of participants. In total, 131 nodes were present in at least 1 ISN, with 62 (47.3%) of those nodes occurring in ≤2 participants (**Supplementary Table 1**). These results demonstrate that, while there is a high degree of similarity in activation patterns of this adaptive language task, it is still important to account for interindividual variability in the organization and precise localization of the language network.

**Figure 2.**
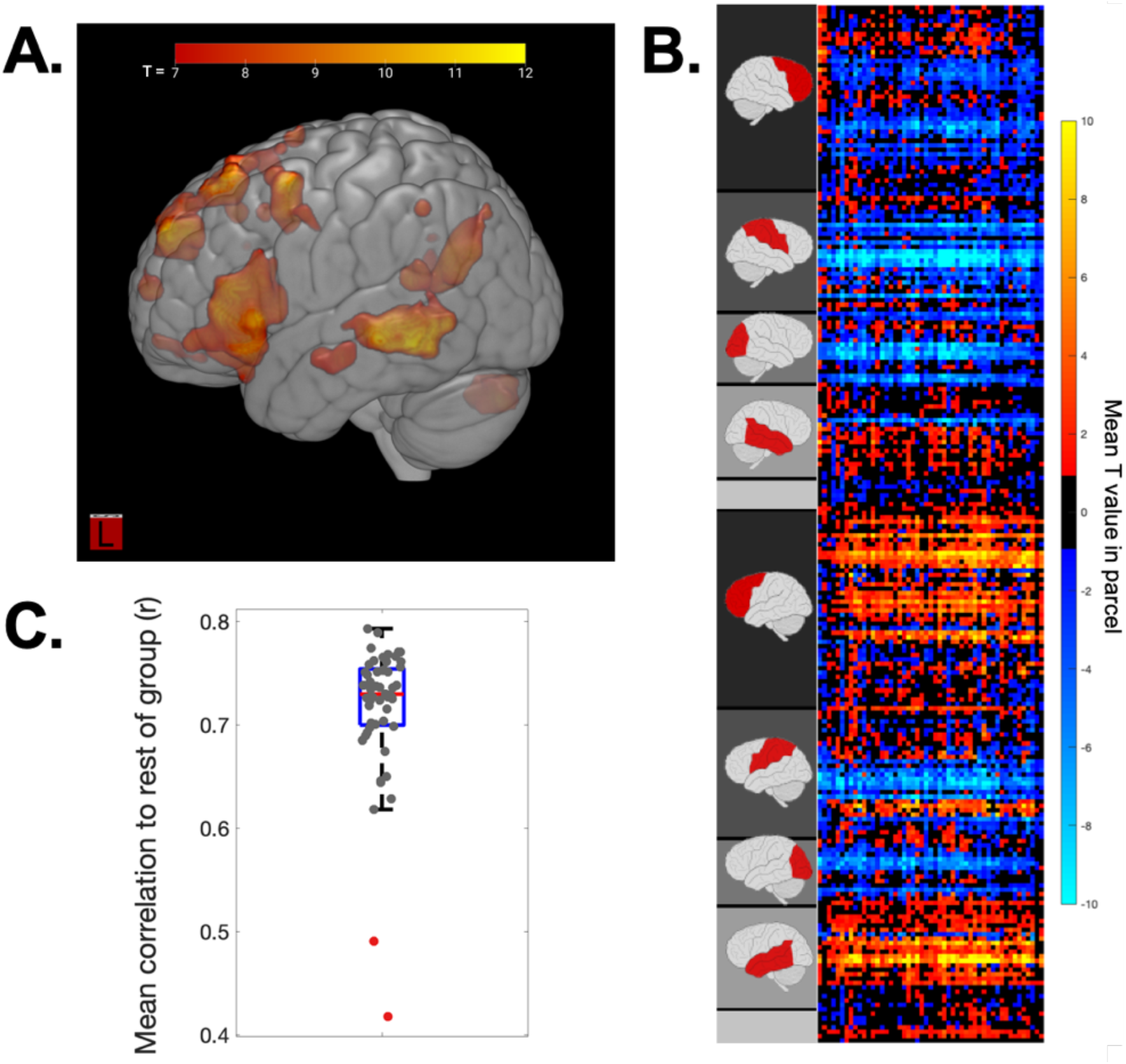
Consistency of task activation for defining semantic network. **A**. Mean voxelwise activation projected onto a standardized brain. Here we show the mean T-values across participants rather than group-level statistics because we are interested in activation at the individual level. **B**. Nodewise activation in all 53 participants (T-values). Each row represents 1 node and each column represents 1 participant. Nodes are sorted by brain lobe according to labels at left (gray box is subcortical). While there are areas of consistent activation in left frontal, temporal, and parietal lobe, there is some interindividual variability. **C**. Correlation of participants’ activation patterns (Pearson’s r). Correlations were calculated between each pair of participants and averaged to give a mean correlation of each individual to the rest of the group. Two participants were notably different from the rest of the group (r < .5) and were found to be right-lateralized for language (first 2 columns in B.); these participants were excluded from future analyses. Overall correlation between individuals was r = .717 including the 2 right-lateralized individuals, and r = .727 after exclusion (R^2^ = .523).

### Structural hub analysis

A hub region analysis was conducted using SC to examine the influence of individual nodes on the rest of the network. In total, 29 hub regions were identified, of which only 2 were present in at least 15 participants (**Figure 3** and **Supplementary Table 2**), with no other nodes appearing as hubs in more than 5 participants. Both of the reliable hub nodes fell within left IFG, specifically in pars triangularis and opercularis, with at least one of these nodes identified as a hub in 33/51 participants. Notably, the pars triangularis node was the only node that was present in 100% of participants’ ISNs. The average participation coefficient of the pars triangularis node was 0.217 ± 0.092, consistent with the definition of a provincial (within-network) node. The other hub in pars opercularis had an average participation coefficient of 0.283 ± 0.111, on the border of a provincial and connector (cross-network) hub. This pars opercularis node was present in 74.5% of participants’ ISNs and was identified as a hub in 42% of these networks.

**Figure 3.**
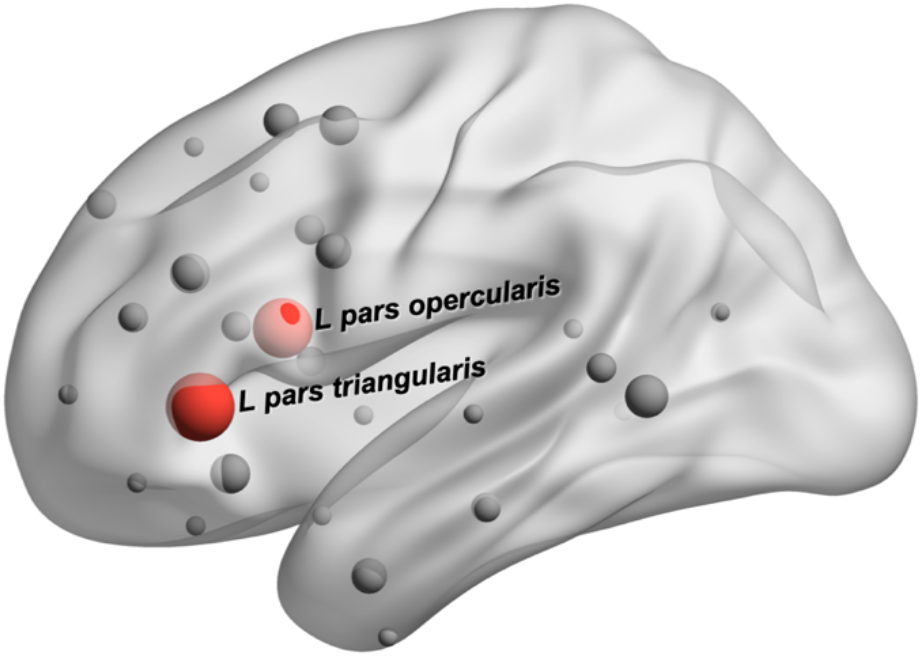
Structural hubs in the semantic network. The size of nodes depicts the number of participants in whom the node was a hub. Nodes in red were found to be hubs in at least 15 participants. For a full list of all nodes identified as hubs, see **Supplementary Table 2**.

### Structural modularity analyses

To interrogate the arrangement of the possible anatomical routes for information to take through the language network, modularity analyses were conducted using structural connectivity. ISNs contained between 3 and 6 modules (mean = 4.27 ± 0.83). Consensus clustering was then performed to enable comparisons across individuals. This second-level analysis revealed 3 consensus modules shown in **Figure 4** and fully detailed in **Supplementary Table 1**. Expected language regions were primarily contained in Modules 2 (green) and 3 (blue). Module 2 covered a swathe of lateral frontal cortex, extending dorsally from left IFG to premotor area and somatosensory cortex. Module3 comprised most of left temporoparietal cortex, extending from temporal pole to the inferior parietal lobule and including anterior parts of left fusiform gyrus. Module1 (red) was more variable across individuals, consistently including default mode regions (Raichle, 2015) in dorsomedial prefrontal cortex, medial orbitofrontal cortex, and parts of posterior cingulate, and less consistently including right-hemisphere homotopes of canonical language regions in temporoparietal cortex and inferior frontal gyrus.

**Figure 4.**
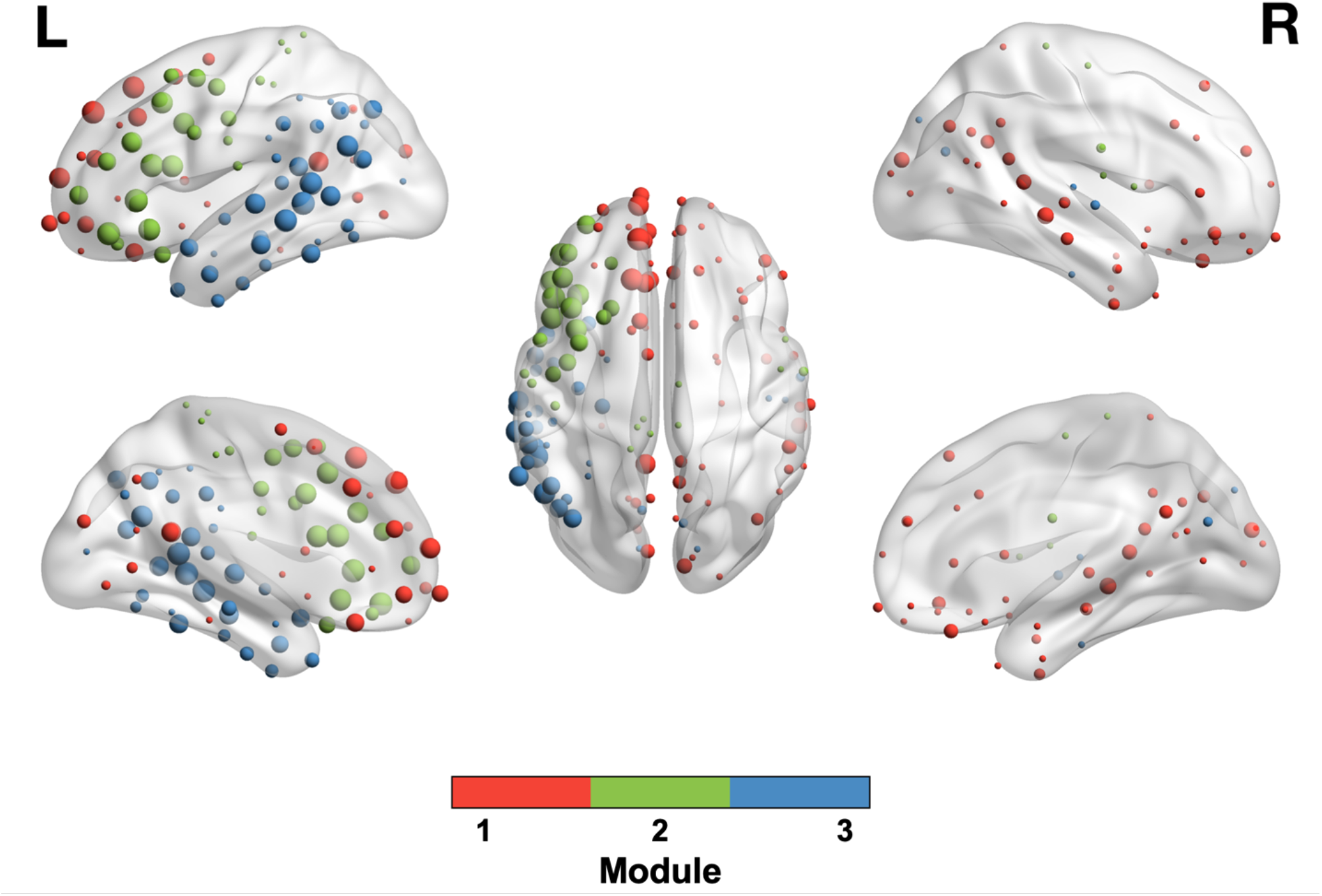
Consensus modules visualized on a model brain. Nodes are color-coded according to module assignment. Node size is proportional to frequency of occurrence of that node in ISNs. Full details are available in **Supplementary Table 1**. The top row shows lateral view; the bottom row shows medial view; and the middle image shows dorsal view.

### Functional connectivity analyses

#### Segregation and integration of the whole network

At the whole-network level, segregation and integration properties were summarized using clustering and global efficiency, respectively, as well as small-world propensity (SWP), a combined measure. Mean values for these measures were as follows: clustering 0.218 ± 0.0513; global efficiency 0.299 ± 0.0674; SWP 0.5036 ± 0.138. In comparison to null networks generated as part of the SWP calculation (Muldoon et al., 2016), ISWs were near ideal levels of clustering (Δ_C_ = 0 for 50/51 participants, where 0 is optimal, i.e. clustered similar to a lattice graph) while path length (efficiency) was less optimized (mean Δ_L_ = 0.70 ± 0.20, where 1 is optimal, i.e. as efficient as a random graph). It has been previously reported that structural and functional brain networks change over the typical aging process (Damoiseaux, 2017) such that they become less segregated and more integrated, i.e., less strongly connected within a network and more strongly connected across networks. To test for this relationship in the current data, correlations (two-tailed Spearman’s rho) were calculated between age and all connectivity measures. No significant correlations were found, suggesting that the connectivity of the semantic network may not be affected by typical aging in the same way as other networks (Chan et al., 2014; Damoiseaux, 2017).

### Segregation and integration of consensus modules

After obtaining the consensus clusters, functional connectivity was calculated within and between modules in each ISN (**Figure 5**). The ISN of one participant with bilateral activation contained only 1 node in Module2 and therefore within-Module2 connectivity could not be calculated; this participant was excluded from all modularity analyses (final *N*=50). Two separate one-way ANOVAs were conducted to test if within-module FC was higher than cross-module FC, as expected, and to explore if connections within or across specific modules were stronger or weaker than the rest of the network. In the first ANOVA, the FC of each edge from each ISN was coded in a binary way as either within-module (i.e., connecting two nodes in the same module) or cross-module (i.e., connecting two nodes in different modules), irrespective of the specific module the nodes were in. FC of within-module edges was significantly higher than cross-module edges (F(1,303))=144.99, P < .0001). In the second ANOVA, edges were coded as one of 6 edge types based on the specific module(s) being linked by the edge, e.g. within-Module1 or Module1-Module2. There was a significant main effect of edge type (F(5,299)=35.61, *P* < .001).

**Figure 5.**
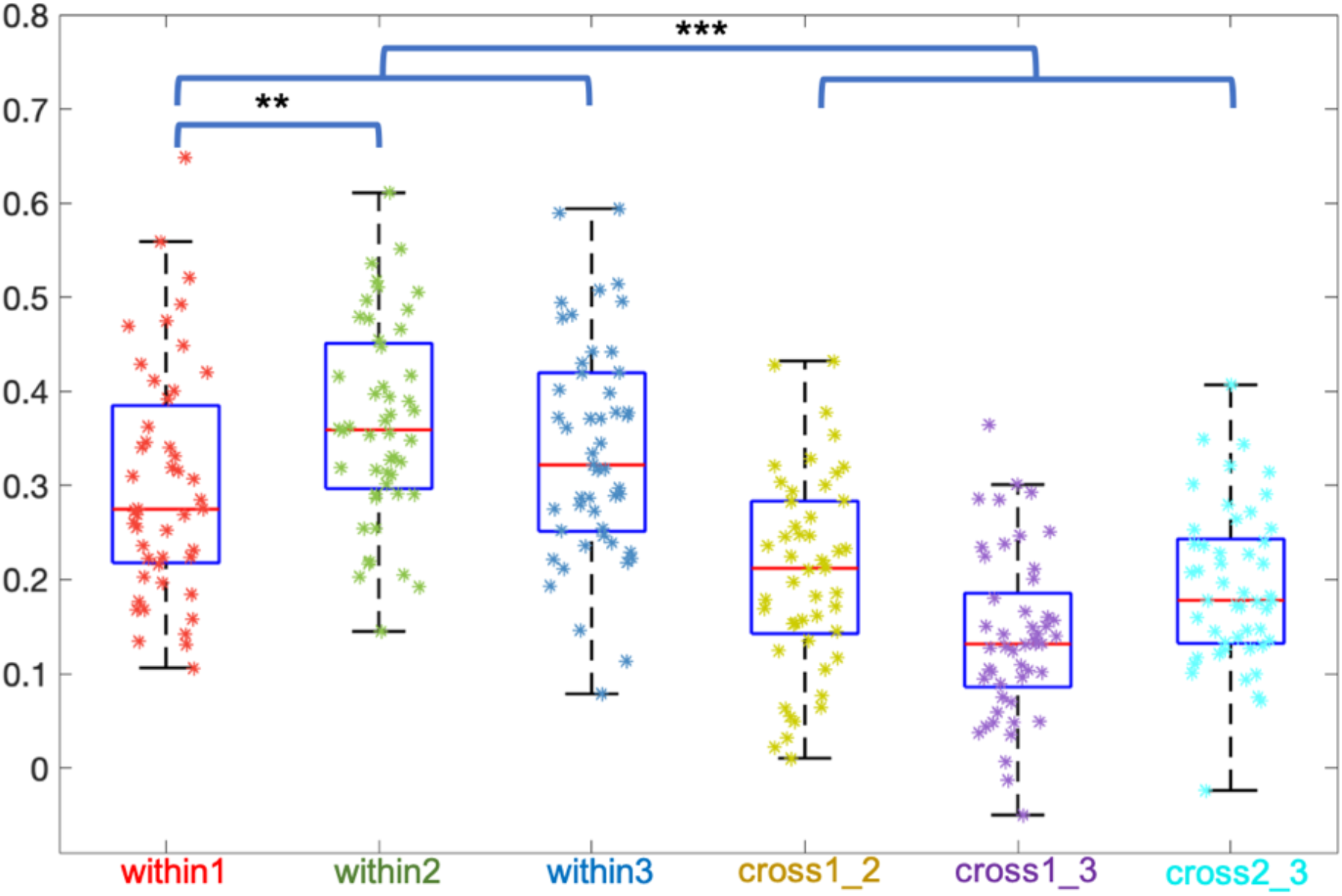
Average functional connectivity within and across structurally-derived consensus modules. FC within modules was greater than connectivity across modules (P < .001), and within-module1 connectivity was lower than within-module2 (P < .01). ^**^ P < .01; ^***^ P < .001

Pairwise post-hoc tests conducted with Tukey’s method showed that FC within Module2 (green) was higher than FC within Module1 (red) (95% CI=[-0.133 -0.012], *P* < .01), but FC in Module3 (blue) was not significantly different from the other two (vs. Module1: 95% CI [-0.105 0.015], *P* = .27; vs. Module2: 95% CI [-0.033 0.088], *P* = .79). Additionally, cross-module FC between Modules 1 and 2 (“Module1×2”) was significantly higher than Module1×3 (95% CI [0.009 0.129], *P* = .014). Module2×3 FC was not significantly different from Module1×2 or 1×3 (vs Module1×2: 95% CI [-0.045 0.075], *P* = .98; vs. Module1×3, 95% CI [-0.114 -0.054]: *P* = .11). In summary, Module2 was highly segregated with strong integration with both Modules 1 and 3; Module1 was not highly segregated; and Module3 seems to be between Modules 1 & 2 on both segregation (FC within modules) and integration (FC across modules).

### Regressions with behavior

To test the hypothesis that verbal fluency in neurotypical adults is related to semantic network organization, backward-elimination linear regressions were performed. These regressions included performance on Letter Fluency and Category Fluency tasks as dependent variables; measures of segregation and integration as predictors; and age and education as covariates. It was hypothesized that semantic function was related to information transfer through the network, so positive relationships were expected with whole-network integration (clustering). For modular connectivity, it was expected that semantic performance would be related to FC within Module3 (temporoparietal cortex), which contains the putative hub endorsed by the hub-and-spoke model, and/or FC between Module2 (left inferior frontal/premotor cortex) and Module3, the proposed pathway in the CSC model of semantic function. Relationships were not expected with Letter Fluency, since the localizer fMRI task was semantic in nature, and the networks being studied were individualized to account for idiosyncrasies in participants’ semantic networks. Two regression models were run. Model #1 featured the whole-network graph theory measures (clustering and global efficiency) as predictors. Model #2 featured within- and cross-module FC. No relationships were found with Letter Fluency (final Model #1: F(1,49)=0.765, *P* = .386; final Model #2: F(1,48)=0.954, *P* = .334).

For Category Fluency, Model #1 demonstrated a positive relationship with education (*P* = .015) and global efficiency (*P* = .003), and a negative relationship with clustering (*P* = .010; overall final Model #1: F(3,47) = 4.499, *P* = .007, adjusted R^2^ = .174; **Table 1**). For the module-based analysis, task performance was positively related to within-Module3 FC (*P* = .031) and marginally related to education (*P* = .060; overall final Model #2: F(2,47)=3.940, *P* = .026, adjusted R^2^ = .107; **Table 1**).

**Table 1.**
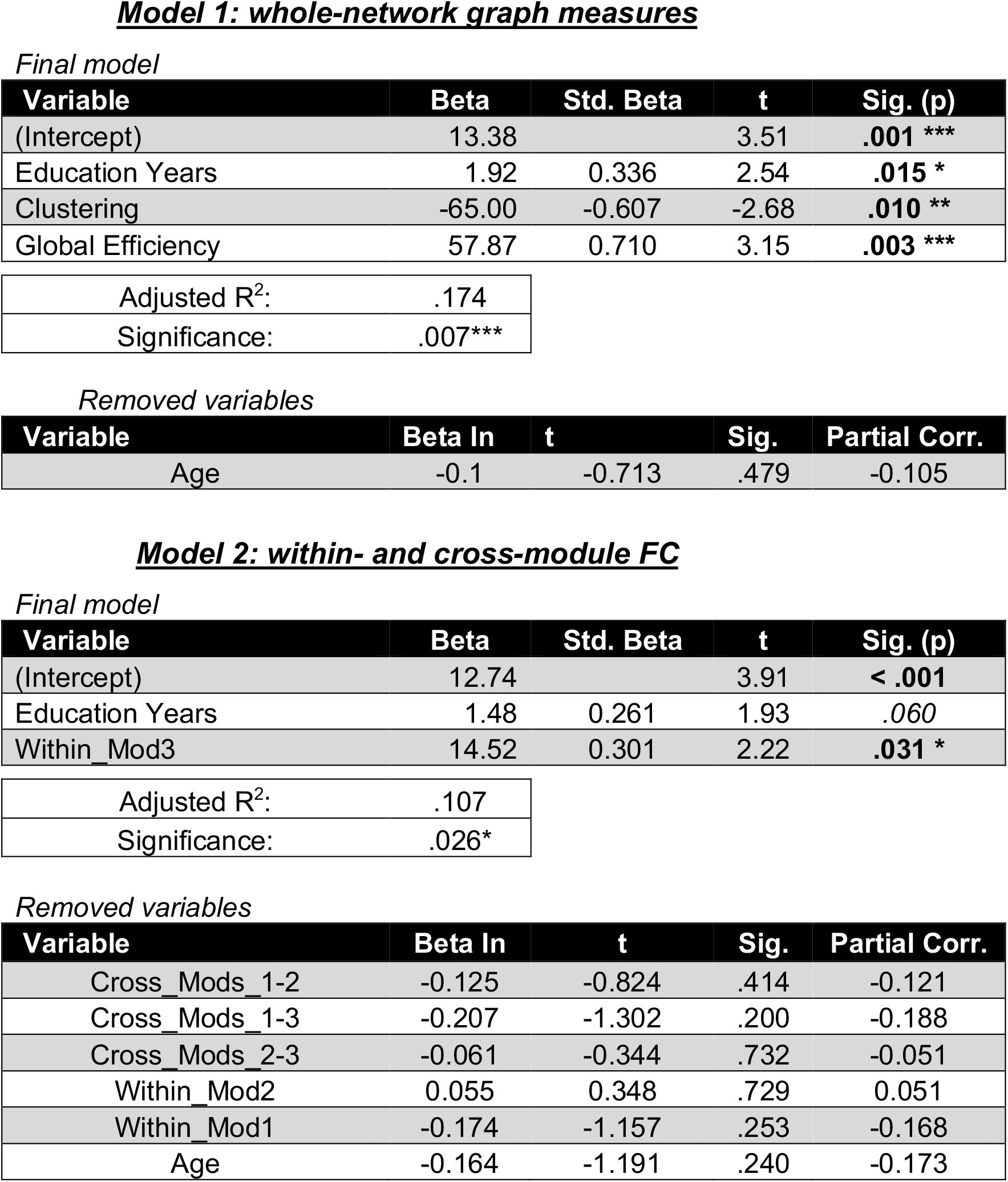
Backward-elimination linear regressions relating performance on Category Fluency, a semantic fluency task, to functional connectivity-derived properties of the semantic network. The final models are shown along with the variables removed by the elimination procedure. ^*^ P < .05; ^**^ P <= .01; ^***^ P < .001

*Final model*
| Variable | Beta | Std. Beta | t | Sig. (p) |
| --- | --- | --- | --- | --- |
| (Intercept) | 13.38 |  | 3.51 | .001 *** |
| Education Years | 1.92 | 0.336 | 2.54 | .015 * |
| Clustering | -65.00 | -0.607 | -2.68 | .010 ** |
| Global Efficiency | 57.87 | 0.710 | 3.15 | .003 *** |
| Adjusted R<sup>2</sup>: | .174 |  |  |  |
| Significance: | .007*** |  |  |  |

*Removed variables*
| Variable | Beta In | t | Sig. | Partial Corr. |
| --- | --- | --- | --- | --- |
| Age | -0.1 | -0.713 | .479 | -0.105 |

**Model 2: within- and cross-module FC** *Final model*
| Variable | Beta | Std. Beta | t | Sig. (p) |
| --- | --- | --- | --- | --- |
| (Intercept) | 12.74 |  | 3.91 | < .001 |
| Education Years | 1.48 | 0.261 | 1.93 | .060 |
| Within_Mod3 | 14.52 | 0.301 | 2.22 | .031 * |
| Adjusted R<sup>2</sup>: | .107 |  |  |  |
| Significance: | .026* |  |  |  |

*Removed variables*
| Variable | Beta In | t | Sig. | Partial Corr. |
| --- | --- | --- | --- | --- |
| Cross_Mods_1-2 | -0.125 | -0.824 | .414 | -0.121 |
| Cross_Mods_1-3 | -0.207 | -1.302 | .200 | -0.188 |
| Cross_Mods_2-3 | -0.061 | -0.344 | .732 | -0.051 |
| Within_Mod2 | 0.055 | 0.348 | .729 | 0.051 |
| Within_Mod1 | -0.174 | -1.157 | .253 | -0.168 |
| Age | -0.164 | -1.191 | .240 | -0.173 |

## Discussion

The goal of this paper was to characterize the structural and functional properties of the individually-defined semantic network and their relation to behavior in a cohort of typical older adults. Studying the organization of the semantic network in typically-aging adults is imperative to gain a better understanding of the brain changes that underpin semantic deficits and recovery in post-stroke aphasia. Semantic network individualization revealed areas of consistent activation during the fMRI task, but also a high degree of interindividual variability in activation pattern.

The hub region analyses identified hubs in left inferior frontal gyrus, most consistently in pars triangularis. The structural modularity analysis revealed three modules in the semantic network, two which align well with canonical left-hemisphere language regions and one default mode/right hemisphere module. We found that performance on a semantic fluency task – but not a phonological fluency task – was related to connectivity within the left temporoparietal module and the segregation and integration of the network as a whole. Together, these results demonstrate an integrated structurofunctional semantic network comprised of subnetworks which may play more important roles for specific processes.

### Hub analyses

Hub regions are thought to be important for coordinating information transfer through brain networks (van den Heuvel & Sporns, 2013b). In our structural hub analysis, we identified only 2 nodes with consistent hub characteristics across individuals, pars triangularis and pars opercularis in left IFG. Notably, pars triangularis was the most robustly-identified hub and was the only node to appear as part of the ISN in every participant, suggesting a common functional role across individuals. Parts of left IFG, particularly pars triangularis, have been previously described as both a functional and structural hub possibly involved in cognitive control due to its density of connections to medial prefrontal cortex (Milton et al., 2021; Vandenberghe et al., 2013; Xu et al., 2016). Further, rTMS experiments have shown that disruption of anterior IFG is associated with increased latency on semantic but not phonological tasks (Gough et al., 2005; Klaus & Hartwigsen, 2019; Whitney et al., 2010), supporting its role as a critical node in the semantic network.

Notably absent from our hub analysis, however, was left anterior temporal lobe (ATL), which is claimed to contain the central hub for semantic function under the hub-and-spoke model (Patterson & Ralph, 2016). Under this model, multisensory input from perceptual “spokes” is bound together in the ATL hub to create a unified, polymodal semantic representation. FC studies have supported the hub status of the ATL in the semantic network in both clinical populations (Vandenberghe et al., 2013; Zhao et al., 2017) and healthy individuals (Xu et al., 2016; Yu et al., 2018). Structural connectivity analyses have been less consistent in identifying ATL as a highly- and widely-connected hub region, although there seem to be relationships in stroke and semantic dementia between severity of semantic deficit and integrity of ATL-related white matter tracts such as superior and inferior longitudinal fasciculi, inferior fronto-occipital fasciculus, and uncinate fasciculus (Chen et al., 2020; Han et al., 2013; Harvey et al., 2013; Sundqvist et al., 2020; Xing et al., 2017). One possible explanation for this inconsistency is that the ATL “hub” is not a discrete node but a functionally-graded span of cortex (Binney et al., 2012; Ralph et al., 2017), better characterized as a subnetwork of connected processors than a unitary hub region. Areas of high functional convergence within this subnetwork (Bajada et al., 2019) might then be detected in FC studies, which might explain the discrepancy between structural and functional hub analyses. Alternatively, the ATL may function as a hub via indirect connections to sensory networks, which are less likely to be detected by SC methods (Sporns, 2013). In total, our structural hub results support a previously-described modification of the hub-and-spoke model, the controlled semantic cognition (CSC) model (Chiou et al., 2018). The CSC model postulates that cognitive control circuits in left IFG exert influence over the graded hub in ATL in order to select the context-appropriate representation amongst related competitors. The role of the pars triangularis hub region in these cognitive control functions deserves further attention in future studies.

Our analyses may be limited by the selection of only nodes that are activated by a semantic decision task, possibly missing brain-wide hubs by ignoring the connectivity of semantic network nodes to the rest of the brain. Future studies should investigate hubs in functionally-defined networks using other fMRI tasks to localize other language subprocesses (e.g., phonology, syntax), or perform brain-wide structurofunctional hub analyses to determine how hub connectivity relates to language function in healthy individuals.

### Consensus modules

The structural modularity analyses revealed three consensus modules at the group level. Module2 (left frontal cortex from inferior frontal gyrus to premotor area) and Module3 (temporoparietal cortex) constituted the majority of the expected language network, and Module1 (posterior cingulate, medial prefrontal cortex, right-hemisphere language homotopes) included regions that have been associated with language but are not domain-specific (i.e., they are activated by other cognitive tasks as well) (Campbell & Tyler, 2018; Fedorenko, 2014).

These modules align well with a previous FC study of the semantic network (Xu et al., 2016) which also found three modules that were labelled “Default mode network” (DMN), corresponding to our Module1; “Frontoparietal network” (FPN), corresponding primarily to our Module2; and “Perisylvian network” (PSN), corresponding to our Module3. In the present study, structural connectivity was used for the modularity analyses to respect the physical, neurophysiological constraints of brain networks which rely on anatomical connections between regions for communication (Chiou et al., 2018). Therefore, our modules are more spatially contiguous than those described in (Xu et al., 2016). One noticeable difference is the module membership of left inferior parietal nodes, which cluster with left temporal nodes in the present study but were assigned to a module containing left inferior frontal nodes in the prior study.

Additionally, this prior study employed a literature search to define 60 nodes based on standard space coordinates, whereas the present study performed analyses on individually-defined semantic networks which do not necessarily include the same nodes across participants. Using a functional localizer at the individual level can reveal idiosyncrasies that are missed by group-level analyses (Gordon et al., 2017). The present results reveal a much broader semantic network than the one reported previously, including more right hemisphere regions which possibly reflects interindividual variability in language network organization (Fedorenko et al., 2010).

### Relationships with behavior

Regressions showed relationships between properties of the semantic network and performance on Category Fluency, a semantic task, but not Letter Fluency, a phonological task. In terms of whole network measures, Category Fluency performance was positively related to integration (efficiency) and negatively related to segregation (clustering). Examining integration and segregation of the network modules, Category Fluency performance was positively related to FC within Module 3 (temporoparietal cortex). It is worth noting that the variance in behavior explained by these models was relatively low, so work remains to identify network properties that robustly explain language ability. Still, these results suggest that functional integration of the semantic network – that is, efficient communication across the whole network – is important for building and accessing semantic representations.

The finding that Category Fluency is positively related to FC within Module3 supports previous results and models of semantic function. These results fit well with the CSC model described earlier in that Module3 would contain the putative graded hub in anterior temporal lobe, and stronger FC within this subnetwork could support access to semantic representations. The results also align with a previous study of healthy native Chinese speakers, which found that semantic performance on reading tasks was positively related to within-language network FC (similar to efficiency) of posterior superior temporal gyrus (pSTG). Follow-up analysis revealed that connections between pSTG, anterior temporal lobe, and anterior fusiform gyrus were most important for semantic function (Yu et al., 2018). Notably, Module3 in the current study contained all of these regions, so our results support this prior finding.

Also of note, this previous study found relationships between phonological performance and language network connectivity to anterior temporal lobe (Yu et al., 2018), whereas we did not find relationships with our phonologically-related task (Letter Fluency). The key difference between this prior study and the current study was the method for identifying network nodes; the prior study selected the same atlas regions for every participant, while the present study identified ISNs using a semantic fMRI localizer. Thus, since our individualized networks were defined using a semantic task, we would not expect to find relationships with phonology. The difference in behavioral results for phonological performance between these studies might reflect the utility of the individual localizer task to identify person-specific nodes involved in a specific process of interest. Future connectivity studies should use multiple localizer tasks designed to reliably parse out brain regions involved in specific language subprocesses to examine how the broader language network dynamically responds across conditions (Fedorenko & Thompson-Schill, 2014).

## Conclusion

In this study, we used multimodal neuroimaging to investigate the properties of the individually-localized semantic network. We used structural connectivity to identify hub nodes and probe the modular organization of the semantic network, revealing 2 hubs in left IFG and a three-module arrangement. While we did not find a structural hub node in ATL as predicted by the hub-and-spoke model (Patterson & Ralph, 2016), we found evidence supporting the importance of within-temporoparietal FC in performance of a semantic fluency task, consistent with previous studies (Yu et al., 2018) and supporting the controlled semantic cognition model (Chiou et al., 2018). Future studies should further investigate the behavioral relevance of these semantic network modules and hub regions to improve our understanding of conditions like semantic aphasia and semantic dementia.

