## Supplementary material for "The organization of individually mapped structural and functional semantic networks in aging adults": Figure S1

### SUPPLEMENTARY FIGURES

#### Graph theory & small-world networks

Graph theory provides mathematical tools for measuring and comparing networks. Prior work has demonstrated that brain networks have the following properties, meeting the definition of a “small-world” network, illustrated on the toy network at right:

- Modularity: presence of communities, or modules, which are more strongly connected within a module than between modules
- Segregation: a high level of interconnectivity among neighboring nodes
  - Often measured with clustering, the probability that, if node X is connected to node Y, node X is also connected to neighbors of node Y. Can be conceptualized by counting triangles in the network
- Integration: a relatively small number of long-distance connections bridging local communities that greatly reduce the distance connecting any 2 nodes
  - Often measured with shortest path length (SPL), the sum of edges required to connect node pairs (illustrated from node X to node Z). Also commonly measured using global efficiency, the inverse of the average SPL for an entire network (high efficiency = short path lengths)
- Hubs are disproportionately highly connected nodes, generally classified as either 1) cross-community “connector” hubs (node X); or 2) within-community “provincial” hubs (node Z)
  - Can be quantified with centrality, a family of measures that describe various aspects of a node’s prominence in a network. Betweenness centrality is the number of shortest paths in the network that pass through a given node; all paths across modules in the toy model must pass through node Z, giving it high betweenness

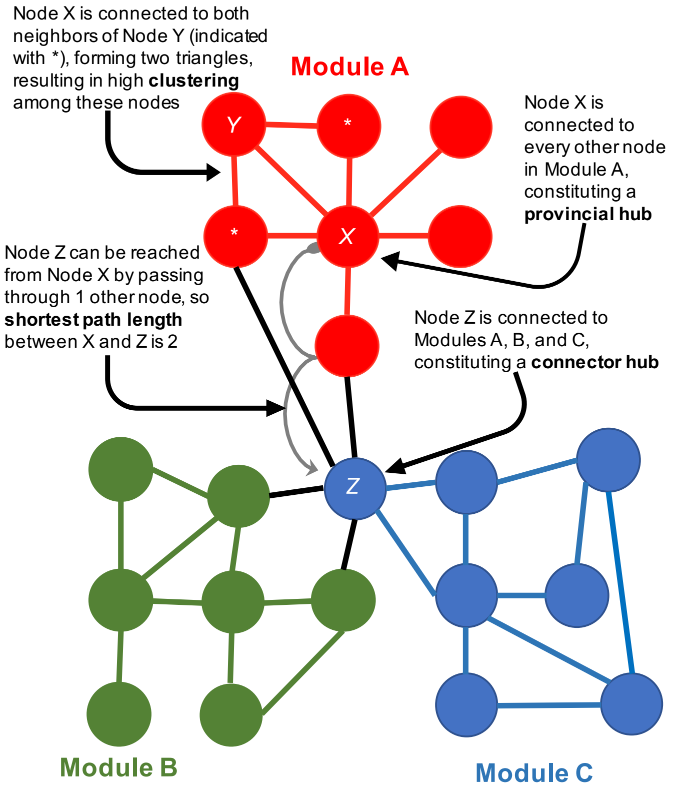


*Figure S1 – Summary of small-world network properties and relevant graph theory measures*
