## Supplementary Table 1 for "The organization of individually mapped structural and functional semantic networks in aging adults"

### SUPPLEMENTARY TABLES

*Table S1 – Full results of consensus clustering showing the name and ID# (nodeid) of each of the nodes from the Lausanne scale 125 parcellation (Hagmann et al., 2008) assigned to the three modules and the count of how many participants' ISNs included each node (count). Module1 was the largest module with 65 nodes; Module2 was the smallest with 29 nodes; and Module3 contained 38 nodes.*

#### Module1

| # | nodeid | name | count |
|---|---|---|---|
| 1 | 135 | L_superiorfrontal_3 | 44 |
| 2 | 137 | L_superiorfrontal_5 | 38 |
| 3 | 133 | L_superiorfrontal_1 | 28 |
| 4 | 159 | L_isthmuscingulate_1 | 24 |
| 5 | 136 | L_superiorfrontal_4 | 18 |
| 6 | 123 | L_medialorbitofrontal_2 | 16 |
| 7 | 134 | L_superiorfrontal_2 | 14 |
| 8 | 122 | L_medialorbitofrontal_1 | 12 |
| 9 | 121 | L_frontalpole_1 | 11 |
| 10 | 139 | L_superiorfrontal_7 | 10 |
| 11 | 96 | R_middletemporal_2 | 10 |
| 12 | 73 | R_cuneus_2 | 8 |
| 13 | 99 | R_bankssts_1 | 7 |
| 14 | 189 | L_cuneus_1 | 6 |
| 15 | 8 | R_medialorbitofrontal_2 | 6 |
| 16 | 68 | R_precuneus_2 | 5 |
| 17 | 5 | R_parsorbitalis_1 | 5 |
| 18 | 140 | L_superiorfrontal_8 | 4 |
| 19 | 132 | L_rostralmiddlefrontal_6 | 4 |
| 20 | 97 | R_middletemporal_3 | 4 |
| 21 | 64 | R_inferiorparietal_4 | 4 |
| 22 | 44 | R_isthmuscingulate_1 | 4 |
| 23 | 198 | L_lingual_3 | 3 |
| 24 | 101 | R_superiortemporal_2 | 3 |
| 25 | 91 | R_inferiortemporal_1 | 3 |
| 26 | 23 | R_superiorfrontal_4 | 3 |
| 27 | 21 | R_superiorfrontal_2 | 3 |
| 28 | 228 | L_caudate | 2 |
| 29 | 197 | L_lingual_2 | 2 |
| 30 | 186 | L_precuneus_3 | 2 |
| 31 | 130 | L_rostralmiddlefrontal_4 | 2 |
| 32 | 110 | R_caudate | 2 |
| 33 | 98 | R_middletemporal_4 | 2 |
| 34 | 69 | R_precuneus_3 | 2 |
| 35 | 62 | R_inferiorparietal_2 | 2 |
| 36 | 29 | R_caudalmiddlefrontal_2 | 2 |
| 37 | 11 | R_parstriangularis_2 | 2 |
| 38 | 6 | R_frontalpole_1 | 2 |
| 39 | 231 | L_accumbensarea | 1 |
| 40 | 229 | L_putamen | 1 |
| 41 | 204 | L_parahippocampal_1 | 1 |
| 42 | 188 | L_precuneus_5 | 1 |
| 43 | 155 | L_rostralanteriorcingulate_1 | 1 |
| 44 | 128 | L_rostralmiddlefrontal_2 | 1 |
| 45 | 119 | L_lateralorbitofrontal_4 | 1 |
| 46 | 115 | R_amygdala | 1 |
| 47 | 114 | R_hyppocampus | 1 |
| 48 | 113 | R_accumbensarea | 1 |
| 49 | 104 | R_superiortemporal_5 | 1 |
| 50 | 95 | R_middletemporal_1 | 1 |
| 51 | 90 | R_temporalpole_1 | 1 |
| 52 | 89 | R_entorhinal_1 | 1 |
| 53 | 82 | R_lingual_2 | 1 |
| 54 | 67 | R_precuneus_1 | 1 |
| 55 | 65 | R_inferiorparietal_5 | 1 |
| 56 | 55 | R_superiorparietal_2 | 1 |
| 57 | 18 | R_rostralmiddlefrontal_5 | 1 |
| 58 | 15 | R_rostralmiddlefrontal_2 | 1 |
| 59 | 9 | R_medialorbitofrontal_3 | 1 |
| 60 | 7 | R_medialorbitofrontal_1 | 1 |
| 61 | 4 | R_lateralorbitofrontal_4 | 1 |
| 62 | 3 | R_lateralorbitofrontal_3 | 1 |
| 63 | 2 | R_lateralorbitofrontal_2 | 1 |
| 64 | 76 | R_lateraloccipital_1 | 1 |
| 65 | 75 | R_pericalcarine_2 | 1 |

#### Module2

| # | nodeid | name | count |
|---|---|---|---|
| 1 | 124 | L_parstriangularis_1 | 51 |
| 2 | 125 | L_parsopercularis_1 | 50 |
| 3 | 120 | L_parsorbitalis_1 | 48 |
| 4 | 126 | L_parsopercularis_2 | 38 |
| 5 | 117 | L_lateralorbitofrontal_2 | 38 |
| 6 | 142 | L_caudalmiddlefrontal_1 | 35 |
| 7 | 143 | L_caudalmiddlefrontal_2 | 31 |
| 8 | 127 | L_rostralmiddlefrontal_1 | 29 |

| # | nodeid | name | count |
|---|---|---|---|
| 9 | 131 | L_rostralmiddlefrontal_5 | 18 |
| 10 | 144 | L_caudalmiddlefrontal_3 | 16 |
| 11 | 129 | L_rostralmiddlefrontal_3 | 16 |
| 12 | 149 | L_precentral_5 | 13 |
| 13 | 116 | L_lateralorbitofrontal_1 | 11 |
| 14 | 150 | L_precentral_6 | 9 |
| 15 | 138 | L_superiorfrontal_6 | 8 |
| 16 | 151 | L_precentral_7 | 7 |
| 17 | 118 | L_lateralorbitofrontal_3 | 5 |
| 18 | 166 | L_postcentral_7 | 2 |
| 19 | 165 | L_postcentral_6 | 2 |
| 20 | 163 | L_postcentral_4 | 2 |
| 21 | 46 | R_postcentral_2 | 2 |
| 22 | 160 | L_postcentral_1 | 1 |
| 23 | 145 | L_precentral_1 | 1 |
| 24 | 37 | R_paracentral_1 | 1 |
| 25 | 154 | L_paracentral_2 | 1 |
| 26 | 153 | L_paracentral_1 | 1 |
| 27 | 45 | R_postcentral_1 | 1 |
| 28 | 38 | R_paracentral_2 | 1 |
| 29 | 31 | R_precentral_1 | 1 |

### Module3

| # | nodeid | name | count |
|---|---|---|---|
| 1 | 216 | L_bankssts_2 | 50 |
| 2 | 215 | L_bankssts_1 | 45 |
| 3 | 212 | L_middletemporal_2 | 44 |
| 4 | 181 | L_inferiorparietal_3 | 34 |
| 5 | 219 | L_superiortemporal_3 | 25 |
| 6 | 213 | L_middletemporal_3 | 23 |
| 7 | 211 | L_middletemporal_1 | 23 |
| 8 | 217 | L_superiortemporal_1 | 19 |
| 9 | 209 | L_inferiortemporal_3 | 19 |
| 10 | 182 | L_inferiorparietal_4 | 17 |
| 11 | 180 | L_inferiorparietal_2 | 17 |
| 12 | 221 | L_superiortemporal_5 | 16 |
| 13 | 183 | L_inferiorparietal_5 | 15 |
| 14 | 214 | L_middletemporal_4 | 13 |
| 15 | 232 | L_hyppocampus | 8 |
| 16 | 169 | L_supramarginal_3 | 8 |
| 17 | 206 | L_temporalpole_1 | 7 |
| 18 | 218 | L_superiortemporal_2 | 6 |
| 19 | 203 | L_fusiform_4 | 6 |
| 20 | 170 | L_supramarginal_4 | 6 |
| 21 | 210 | L_inferiortemporal_4 | 5 |
| 22 | 207 | L_inferiortemporal_1 | 5 |
| 23 | 220 | L_superiortemporal_4 | 4 |
| 24 | 208 | L_inferiortemporal_2 | 4 |
| 25 | 201 | L_fusiform_2 | 3 |
| 26 | 187 | L_precuneus_4 | 2 |
| 27 | 167 | L_supramarginal_1 | 2 |
| 28 | 102 | R_superiortemporal_3 | 2 |
| 29 | 72 | R_cuneus_1 | 2 |
| 30 | 233 | L_amygdala | 1 |
| 31 | 202 | L_fusiform_3 | 1 |
| 32 | 190 | L_pericalcarine_1 | 1 |
| 33 | 174 | L_superiorparietal_3 | 1 |
| 34 | 171 | L_supramarginal_5 | 1 |
| 35 | 168 | L_supramarginal_2 | 1 |
| 36 | 105 | R_transversetemporal_1 | 1 |
| 37 | 92 | R_inferiortemporal_2 | 1 |
| 38 | 60 | R_superiorparietal_7 | 1 |
