## Supplementary Table 2 for "The organization of individually mapped structural and functional semantic networks in aging adults"

*Table S2 – Summary of the structural hub analysis. All nodes that were identified as hubs in at least 1 participant are listed. ConMod = consensus module; HubCount = number of participants in whom the node was a hub; NodeCount = number of participants whose ISN contained that node; HubPct = HubCount/NodeCount; meanPartCo = average participation coefficient for that node; mean_FC = mean functional connectivity to the entire ISN; FC_modX = mean functional connectivity with module X*

| nodeID | Node name | ConMod | HubCount | NodeCount | HubPct | meanPartCo | mean_FC | FC_mod1 | FC_mod2 | FC_mod3 |
|---|---|---|---|---|---|---|---|---|---|---|
| 124 | L_parstriangularis_1 | 2 | 29 | 51 | 56.9% | 0.217 | 0.281 | 0.201 | 0.425 | 0.233 |
| 126 | L_parsopercularis_2 | 2 | 16 | 38 | 42.1% | 0.283 | 0.300 | 0.185 | 0.415 | 0.288 |
| 211 | L_middletemporal_1 | 3 | 5 | 23 | 21.7% | 0.154 | 0.255 | 0.092 | 0.218 | 0.414 |
| 117 | L_lateralorbitofrontal_2 | 2 | 4 | 38 | 10.5% | 0.184 | 0.226 | 0.207 | 0.308 | 0.169 |
| 127 | L_rostralmiddlefrontal_1 | 2 | 4 | 29 | 13.8% | 0.145 | 0.286 | 0.275 | 0.409 | 0.169 |
| 144 | L_caudalmiddlefrontal_3 | 2 | 4 | 16 | 25.0% | 0.193 | 0.316 | 0.322 | 0.418 | 0.208 |
| 138 | L_superiorfrontal_6 | 2 | 3 | 8 | 37.5% | 0.372 | 0.330 | 0.403 | 0.392 | 0.194 |
| 151 | L_precentral_7 | 2 | 3 | 7 | 42.9% | 0.192 | 0.267 | 0.137 | 0.382 | 0.194 |
| 214 | L_middletemporal_4 | 3 | 3 | 13 | 23.1% | 0.071 | 0.199 | 0.176 | 0.156 | 0.264 |
| 125 | L_parsopercularis_1 | 2 | 2 | 50 | 4.0% | 0.151 | 0.288 | 0.187 | 0.416 | 0.257 |
| 129 | L_rostralmiddlefrontal_3 | 2 | 2 | 16 | 12.5% | 0.248 | 0.220 | 0.227 | 0.369 | 0.051 |
| 135 | L_superiorfrontal_3 | 1 | 2 | 44 | 4.5% | 0.312 | 0.263 | 0.370 | 0.295 | 0.174 |
| 143 | L_caudalmiddlefrontal_2 | 2 | 2 | 31 | 6.5% | 0.161 | 0.317 | 0.241 | 0.412 | 0.277 |
| 213 | L_middletemporal_3 | 3 | 2 | 23 | 8.7% | 0.089 | 0.270 | 0.195 | 0.201 | 0.439 |
| 215 | L_bankssts_1 | 3 | 2 | 45 | 4.4% | 0.049 | 0.258 | 0.115 | 0.190 | 0.460 |
| 228 | L_caudate | 1 | 2 | 2 | 100.0% | 0.460 | 0.108 | 0.163 | 0.135 | 0.032 |
| 95 | R_middletemporal_1 | 1 | 1 | 1 | 100.0% | 0.008 | 0.342 | 0.415 | 0.084 | 0.223 |
| 120 | L_parsorbitalis_1 | 2 | 1 | 48 | 2.1% | 0.226 | 0.255 | 0.234 | 0.354 | 0.195 |
| 123 | L_medialorbitofrontal_2 | 1 | 1 | 16 | 6.3% | 0.349 | 0.119 | 0.201 | 0.126 | 0.073 |
| 131 | L_rostralmiddlefrontal_5 | 2 | 1 | 18 | 5.6% | 0.392 | 0.215 | 0.306 | 0.299 | 0.066 |
| 137 | L_superiorfrontal_5 | 1 | 1 | 38 | 2.6% | 0.485 | 0.277 | 0.389 | 0.325 | 0.162 |
| 142 | L_caudalmiddlefrontal_1 | 2 | 1 | 35 | 2.9% | 0.380 | 0.296 | 0.345 | 0.378 | 0.196 |
| 180 | L_inferiorparietal_2 | 3 | 1 | 17 | 5.9% | 0.245 | 0.191 | 0.108 | 0.163 | 0.272 |
| 207 | L_inferiortemporal_1 | 3 | 1 | 5 | 20.0% | 0.054 | 0.199 | 0.190 | 0.147 | 0.261 |
| 217 | L_superiortemporal_1 | 3 | 1 | 19 | 5.3% | 0.136 | 0.196 | 0.070 | 0.159 | 0.319 |
| 219 | L_superiortemporal_3 | 3 | 1 | 25 | 4.0% | 0.052 | 0.207 | 0.098 | 0.150 | 0.362 |
| 221 | L_superiortemporal_5 | 3 | 1 | 16 | 6.3% | 0.032 | 0.144 | 0.108 | 0.134 | 0.169 |
| 229 | L_putamen | 1 | 1 | 1 | 100.0% | 0.430 | 0.103 | 0.127 | 0.099 | 0.085 |
